# “SELECTIVE SURVIVAL OF AVIAN ECTOPARASITES IN AGROINTENSIVE HABITATS AND SUBSEQUENT EFFECTS ON HOST SELECTION BY PARASITOIDS”

**DOI:** 10.64898/2026.10.03.756457

**Authors:** Simon Coroller-Chouraki, Jade Savage, Marc Bélisle

## Abstract

Morphological variation in insects can indicate responses to habitat change and land use intensification, with potential consequences for trophic interactions. In insect host–parasitoid systems, host body is often a key determinant of offspring development and emergence success, as well as parasitoid host-selection behavior. Yet, most studies in agricultural landscapes have focused on size variation in crop-associated insects (*i.e.:* pests, pollinators and enemies), leaving those relying on non-crop resources largely unexplored. Here, we investigated how agroenvironmental contexts and trophic resource availability influence puparial size in *Protocalliphora sialia* (Diptera: Calliphoridae), an obligate hematophagous ectoparasite of Tree Swallow (*Tachycineta bicolor*) nestlings. We further tested whether puparial size affects emergence probability and risk of parasitoidism by *Nasonia* spp. (Hymenoptera: Pteromalidae) jewel wasps. Using 1,513 *P. sialia* puparia from 88 nests within a nest-box monitoring system spanning a 10,200-km^2^ east–west intensification gradient in southern Québec, Canada, we evaluated how agrointensive contexts, avian host availability, and ectoparasite densities influence puparial size, parasitoidism risk, and emergence success.

The agricultural context appears to shape puparial size through survival biases and/or altered growth rates. In agroextensive contexts (*i.e.:* pastures, forages with high tree cover), where *Protocalliphora* are more numerous, *P. sialia* puparia were smaller and exhibited greater size variation. Conversely, in agrointensive environments (maize and soybeans monocultures with higher doses of pesticides), where *Protocalliphora* are fewer, small individuals were underrepresented, suggesting a selective mortality in *P. sialia* population. However, puparial size explained less than 10% of the variation in emergence or parasitism, likely reflecting consistently high parasitic pressure from *Nasonia* wasps, which infested 50-90% of puparia, depending on site and year. Our findings highlight the importance of selective mortality in structuring trait distributions and modulating multi-trophic interactions in human-modified landscapes.

## Introduction

Practices associated with intensive agriculture, such as heavy uses of synthetic chemicals (Goulson *et al*. 2015, Uhl and Brühl 2019), frequent machinery use, mowing (Cizek *et al*. 2012, Humbert *et al*. 2009, 2012), and landscape homogenization (Kovács-Hostyánszki *et al*. 2017), impact insect populations and communities both directly and indirectly. These disturbances extend to trophic networks in agroecosystems, reducing floral resources (Goulson *et al*. 2015), disrupting essential interactions between predators and preys (Geiger *et al*. 2010, Isenhour 1985, 1986) or hosts and parasites (Coroller-Chouraki *et al*. 2026c), potentially leading into pest outbreaks (Janssen and van Rijn 2021, Peisley *et al*. 2015, Sánchez-Bayo 2021). Beyond evaluating the impacts of agricultural intensification on insect population dynamics and ecosystem services, research has also focused on its effects on functional responses, behavior, physiology and morphology (Gérard *et al*. 2022, Ponce-Méndez *et al*. 2022, Verheyen and Stoks 2019, Wisniewska and Prokopy 1997).

Body size is a key morphological trait in insect species as it correlates to physiology, life history and ecology (Chown and Gaston 2010, Peters 1986). Variation in body size can reflect the impact of fluctuating environments such as agricultural intensification and land use on species level responses. Intraspecific phenotypic variability in populations can be beneficial (Henry *et al*. 2009, Maynard *et al*. 2019) although reduced size often affects insects foraging ranges, resource use, pollination, reproduction and dispersal abilities (Schweiger et al. 2005, Huang et al. 2016, Beukeboom 2018, Elzay and Baum 2021). Moreover, size reduction can predict both individual condition and subsequent survival (Forrest 2016, Gergs and Jager 2014). Notably, morphological variations can directly affect trophic interactions in cases such as parasitoid-host dynamics where factors like host selection and oviposition are influenced by both host and parasitoid sizes (Henry *et al*. 2009, Kouamé and Mackauer 1991).

Recent studies on bees and bumblebees show a decline in body size under intensive agricultural practices, likely due to increased exposure to pesticide, reduced foraging resources, and reduced nesting opportunities (Elzay and Baum 2021, Gérard *et al*. 2022, Renauld *et al*. 2016). Yet, the number of longitudinal size comparisons conducted across agroecosystems is limited due to labor demands, limited taxonomic expertise, and funding constraints for insect taxa not directly affecting crop production. Moreover, findings remain inconsistent, with strong interspecific variation, and are subject to publication bias, where negative agricultural intensification effects on size are more visible than studies reporting no effect (Gérard *et al*. 2022). A survey focusing more generally on the Diptera family indicates that intensive agricultural practices were associated with diminished total biomass whereas individual counts remain comparable or even higher (Powell *et al*. 2024). These results point to potential community shifts and/or overall size reductions. Such diversity makes order-wide generalizations misleading and underscores the need to focus on specific taxa, trophic niches and networks. Body size can be highly plastic in Calliphoridae, reflecting variation in resource availability and developmental conditions experienced during larval growth (Tarone and Foran 2006).

*Protocalliphora* bird blowflies (Diptera) are obligate hematophagous nestling ectoparasites. While most work emphasizes their costs to avian hosts (Maziarz *et al*. 2022), recent studies have explored ecological drivers of their abundance and infestation (Coroller-Chouraki *et al*. 2026b; Mennerat *et al*. 2021, Merino *et al*. 2023, Musgrave *et al*. 2019, Río *et al*. 2023). Despite growing interest in *Protocalliphora* ecology, research has primarily focused on their effects on avian hosts. Consequently, most ecological studies on bird blowflies have concentrated exclusively on their abundance or prevalence, leaving their potential fitness consequences of morphological plasticity unexplored. To our knowledge, only two publications have investigated *Protocalliphora* puparial size and development, focusing on temperature effects and relying on limited sample sizes under controlled *in-vivo* or *in-vitro* conditions (Bennett and Whitworth 1991, Brouillard 2024). Unlike many phytophagous or canopy-dwelling insect species whose body size variation under agricultural intensification has been measured in individuals developing directly within treated crops (Gérard *et al*. 2022, Renauld *et al*. 2016), measuring *Protocalliphora* puparia provides a different perspective. These ectoparasites larvae develop entirely within bird nests and feed exclusively on the blood of nestling hosts. Consequently, they are not directly exposed to agricultural habitats during development. Any morphological variation associated with the surrounding agricultural landscape must therefore arise through different pathways. These may include host-mediated effects, such as reduced resource availability due to nestling premature death or lower blood quality and higher toxicological load (Sigouin *et al*. 2021), maternally-mediated effects related to the environment experienced by adult females prior to oviposition, or the direct transfer of agricultural contaminants into the nest microhabitat through nesting material, dust deposition, or prey delivered to nestling (Cordi *et al*. 1997, Poisson *et al*. 2021). As agricultural intensification alters *Protocalliphora* infestation rates by *Nasonia* wasps parasitoids (Hymenoptera: Pteromalidae; Coroller-Chouraki *et al*. 2026c), the extent to which landscape-level pressures translate into individual size and subsequent trophic implications remain unknown. Puparial size might influence *Protocalliphora* survival, emergence success, and host selection by parasitoids, making it a potential indicator of agroecosystem disturbance.

Given that agricultural intensification can indirectly ialter ectoparasite development and emergence through host-mediated pathways, the primary objectif of this study was to quantify how both agricultural landscapes and nestling host availability independantly affect *Protocalliphora sialia* puparial size. To do so, we used a 16-year Tree Swallow (*Tachycineta bicolor*) nest-box monitoring program across a gradient of agroenvironmental contexts. Furthermore, we assessed the downstream ecological consequences of this size variation on *P. sialia* emergence success and its susceptibility to parasitoidism by *Nasonia* spp. wasps.

## Materials and Methods

### Geography, agricultural landscape and meteorology

We collected data between 2004 and 2019 from a large-scale nest box network established in southern Québec, Canada, to examine the effects of agricultural land use on aerial insectivores. The network comprises 400 nest boxes evenly distributed among 40 farms spanning a ∼10,200 km^2^ east–west gradient of agricultural intensity (Fig. 1a). Nest boxes were installed ∼1.5 m above ground and spaced ∼50 m apart along field-edge transects. Eastern sites were dominated by pastures, forage, and tree cover, while western sites featured intensive monocultures (soybean, maize) with heavy pesticide use (Giroux 2022). Initially designed to study Tree Swallow (*Tachycineta bicolor*) prey availability (Powell *et al*. 2024) and breeding ecology (Garrett *et al*. 2022), the system has since been used to answer additional questions such as the effects of *Protocalliphora* spp. parasitism on avian reproductive success (Daoust *et al*. 2012a,b, Pigeon *et al*. 2013, Sigouin *et al*. 2021) and parasite responses to agricultural intensification (Coroller-Chouraki *et al*. 2026b).

**Figure 1.**
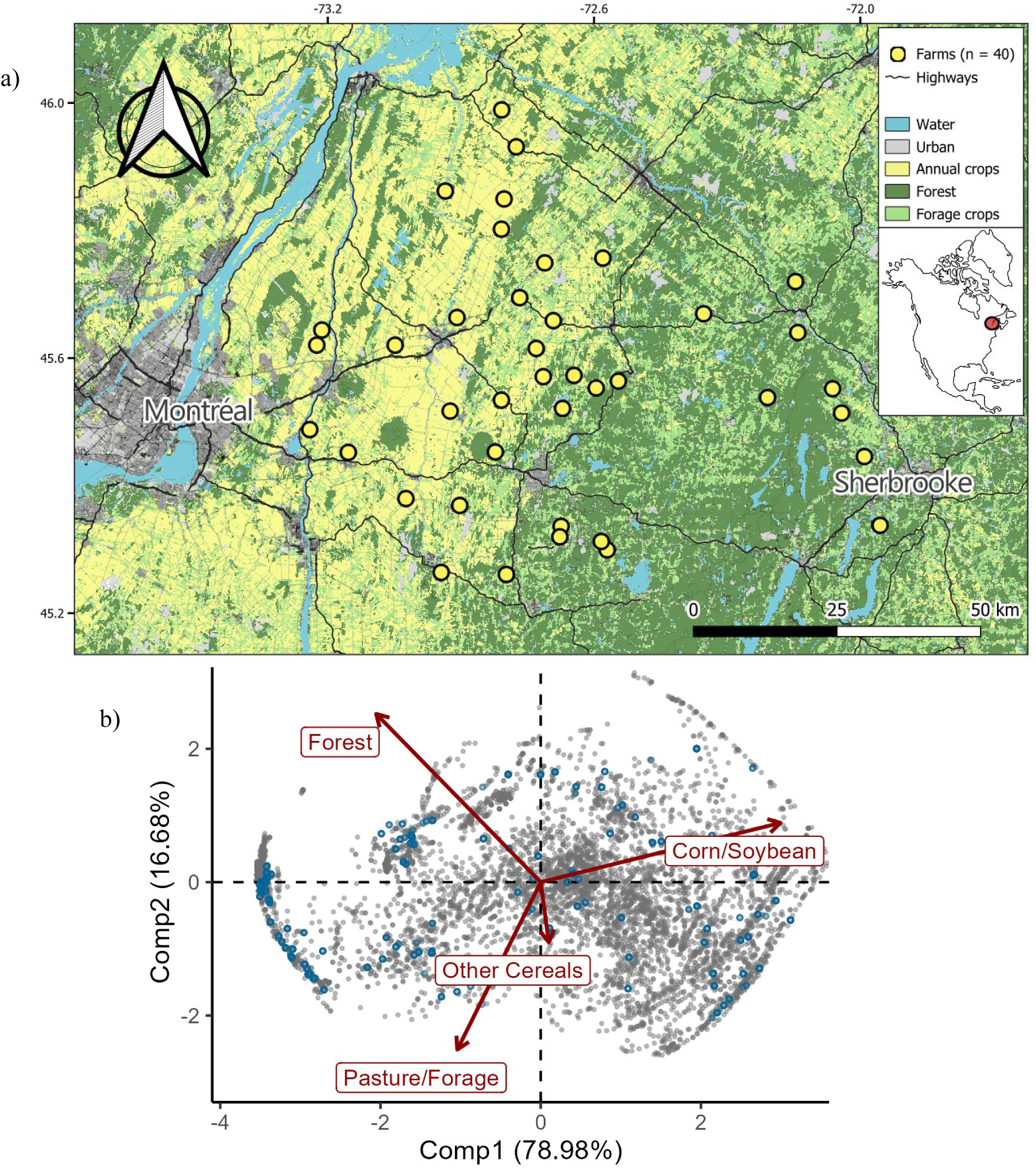
a) Map of the nest-box network composed of 40 farms (10 nest boxes per farm) distributed across a 10,200-km^2^ gradient of agricultural intensification in southern Québec, Canada between 2004 and 2019. Map coordinates follow the WGS 84, EPSG:4326 reference system. Spatial data were projected in the NAD83 MTM Zone 8 system for mapping and spatial analyses. b) biplot of the Robust Principal Components Analysis (rPCA) fitted on the relative land cover values determined annually around each of the 400 nest boxes (500-m radius) of the study system between 2006-2019 (n= 5,600). Blue dots represent the agricultural contexts surrounding the nests where *P. sialia* puparia have been sampled and measured for this study (n=88),

Local landscapes were characterized annually, including crop types in situ around each farm and classified land cover within a 500 m radius of each nest box into four categories: Woodlands, Maize & Soybeans, Other Cereals, and Pasture/Forage/Wasteland. We delineated habitat boundaries from orthophotos (1:40,000) using QGIS and computed proportional cover for each category. We performed a robust compositional PCA (Fig. 1b; Garrett *et al*. 2022) on annual crop and forest cover data collected from 2006 to 2019. The first component (Comp. 1) explained ∼79 % of the variation, representing a gradient from maize/soybean-dominated (positive) to forage/forest-dominated (negative) landscapes. The second component (Comp. 2) explained ∼17 %, contrasting tree cover (positive) with forage cover (negative). Because in situ landscape data were unavailable for 2004–2005, Comp. 1 and Comp. 2 scores for those years were estimated as the mean of the three subsequent years, given the low interannual variation in both axes for the same nest boxes (Fig. S1).

We used the NASAPOWER package in RStudio (v4.3.1; Sparks 2024) to extract hourly temperature (°C) and precipitation (mm) according to the GPS coordinates of each farm at a 50 km resolution. For each nest, meteorological average values were calculated from the first hatching date to seven days after the last Tree Swallow nestling fledged or died, covering the full development and pupation period of *Protocalliphora* spp. (Sabrosky *et al*. 1989).

### Nests and parasites sampling

We monitored Tree Swallows nests every two days to record clutch size, nestling survival, and fledging age. We estimated nestling host availability for *Protocalliphora* as the total number of nestling-days per nest, a key driver of parasite abundance (Coroller-Chouraki *et al*. 2026b). After each breeding season, we collected nests from single clutches with at least one hatched egg. We then sealed, stored at 4 °C, and later froze the nests at –80 °C until processing. We sorted the nest material under a ventilated hood and extracted, counted and preserved *Protocalliphora* puparia in labelled vials containing 75% ethanol. Data were collected from 2004 to 2019, except in 2008–2009 due to methodological inconsistencies (Daoust *et al*. 2012a). During those two years, Daoust *et al*. reared pupae in the laboratory to collect emerging *Protocalliphora* spp. and *Nasonia* spp. wasps, resulting in the loss of specimens for subsequent examinations.

Because the number of puparia recovered from infested nests was exceptionally high over the 14-year period (excluding 2008-2009), a standardized subsampling protocol was implemented to ensure representative coverage of the agricultural gradient while keeping processing time manageable. Subsampling followed this standardized procedure: 10 farms were randomly selected across the study area using a R-based randomization algorithm. Within each selected farm, up to 2 infested nests (when available) were R-based randomly chosen each year, and all puparia they contained were retained. When the total number of puparia for a given year was below 100, additional farms and nests were randomly added following the same procedure until the yearly threshold was reached. For this study we used a total of 88 nests which yielded 1,513 *P. sialia* suitable for morphometric analysis, as well as 1,511 *P. sialia* suitable for parasitoidism and emergence diagnosis. We consolidated these individual and nest-level observations into a comprehensive database, linking each record to the specific agricultural landscape, avian host availability and meteorological vaiables detailed above.

### Species identification and parasitoidism diagnosis

Puparia species identification and the assessment of both emergence and parasitoidism status were performed using keys (Sabrosky *et al*. 1989, Whitworth 2019) and following the protocols detailed in Coroller-Chouraki *et al*. (2026a). Three species were sympatrically present in our study system: *Protocalliphora sialia* (82.6%), *P. bennetti* (15.75%), and *P. metallica* (3.62%). In line with our objectives, we focused exclusively on *P. sialia*, the most abundant species.

As Daoust *et al*. (2012a) reared 956 *Protocalliphora* pupae from the same study system in 2008–2009 and identified two parasitoid species emerging from parasitized individuals (*N. vitripennis* (96.5%) and *N. giraulti* (3.5%)), all emergence holes (0.5–1 mm in diameter) observed in the present dataset were attributed to *Nasonia* spp. For each *P. sialia* puparia examined, evidence of *Nasonia* parasitism (such as exit holes or unemerged *Nasonia* larvae within dissected hosts) was recorded. Failed emergence of *P. sialia* that were not attributable to *Nasonia* parasitoids were recognized by the presence of an intact operculum and pupal remnants without *Nasonia* first instars inside the dissected puparia.

### Morphometric analysis

A total of 1,513 *P. sialia* puparia were photographed with a Leica M165FC stereomicroscope equipped with a 0.63× planApo objective, a Leica MC170HD color camera, and Leica LAS X software (v 3.0.3). The microscope was factory-encoded for pixel-size calibration (µm/pixel), ensuring accurate dimensional measurements. All image-based measurements were performed in Fiji (Schindelin *et al*. 2012) using a custom script developed with the Fiji/ImageJ API. For each image, a threshold-based segmentation isolated individual puparia, and the “Fit Ellipse” function was applied to each object. This approach modelled each puparium as an ellipsoid and calculated its theoretical surface area (mm^2^), which we used as a proxy for size. Measuring this projected area rather than length or width minimizes bias due to missing opercula in emerged puparia, while providing a reproducible 2D metric that reflects both elongation and width variation.

### Statistical analysis

We first built a Directed Acyclic Graph (DAG ; Fig. S2) according to the literature and our knowledge of the ecological system. Such methods allow to fathom the direct, indirect and total effects of one variable on another, as well as controlling for confounding factors (Cinelli *et al*. 2022). As data originated from a hierarchical study design (Nestbox ID, Farm ID, Year), we used mixed models, controlling with these variables as random effects.

Model A (Table 1) was developed to investigate how the surface of *P. sialia* puparia is influenced by agricultural landscape configuration, meteorological conditions during larval development, nestling host availability, and *Protocalliphora* spp. density within the nest (*i.e. :* puparia from all species). For a given number of *Protocalliphora* spp. parasites within the nest, higher nestling availability (providing blood resources and warmth) might create more favorable conditions for larval development, potentially leading to larger individuals. However, hypothetically, this effect might be density-dependent, with increased parasite competition limiting individual growth. To account for these dynamics, an interaction between host availability and parasite density was included in the model. Models B and C (Table 1), explore how the surface of *P. sialia* puparia, when controlling for agroenvironmental parameters, affects the likelihood of infestation by *Nasonia* spp. parasitoid wasps and its emergence success in the absence of parasitoidism. These models incorporating puparial size as an explanatory variable to assess its influence on both infestation probability and emergence outcomes of *P. sialia*.

**Table 1.** Models A), B) C) building according to the Directed Acyclic Graph (Fig. S2). In the case of GAMM analyses, s() refers to smoothing terms, while t2() refers to full-tensor product smooth for interactions. Comp.1 & Comp.2 : Components of the robust Principal Component Analysis.

| Models | Response variable | Regression Family Distribution | Explanatory variables | Random effects Sample size |
| --- | --- | --- | --- | --- |
| A | Body size of *P. sialia* individuals, approximated by puparia's area (mm<sup>2</sup>) & associated variance | GAMM<br>Gaussian<br>(link = "identity") | t2(Nestling availability, #*Protocalliphora* spp. within the nest) + t2(Comp.1,Comp.2) + Mean Temp.+ Mean Precip. + First hatching date | (1\|Year/Farm ID)<br>(1\|Nestbox ID)<br>n = 1,513 |
| B | Individual probability of *Nasonia* spp. infestation on *P. sialia* | GAMM<br>Bernoulli<br>(link = "logit") | *P. sialia* body size (mm<sup>2</sup>) + s(#*Protocalliphora* spp. within the nest) + t2(Comp.1, Comp.2) + Mean Temp. + Mean Precip. + First hatching date | (1\|Year/Farm ID)<br>(1\|Nestbox ID)<br>n = 1,511 |
| C | Emergence probability of *P. sialia* (excluding *Nasonia*-infested ones) | GLMM<br>Bernoulli<br>(link = "logit") | *P. sialia* body size (mm<sup>2</sup>) + #*Protocalliphora* spp. within the nest + t2(Comp.1 x Comp.2) + Mean Temp. + Mean Precip. + Nestling host availability + First hatching date | (1\|Year/Farm ID)<br>(1\|Nestbox ID)<br>n = 305 |

Temp. & Precip.: Temperature and precipitation (averaged during either nestling or *Protocalliphora* development period.

We fitted Generalized Additive Mixed Models (GAMMs) and Generalized Linear Mixed Models (GLMMs) using the *brms* package (v2.21; Bürkner 2021) in R (v.4.3.1, R core teams 2023), specifying non-informative or weakly informative priors. Model predictions were extracted and visualized using the *tidybayes* package (v.3.0.6, Kay 2023). For each model, we plotted the marginal effect of explanatory variables while holding all other variables constant at their means and at the population level. Credibility intervals (50%, 80% and 95%) were highlighted to illustrate uncertainty around the marginal effect estimates.

To compare the predicted morphological variation in *P. sialia* puparial size across contrasting agricultural contexts, we generated posterior predictive distributions under two contrasting landscape scenarios. Scenario 1 (S1; for which Comp.1 = -2 and Comp.2 = 1.3) represented agroextensive farming systems, comprising pastures and forages with high tree cover; scenario 2 (S2; for which Comp.1 = 2.3, Comp.2 = 2.4) represented agrointensive farming systems, comprising maize and soybean monocultures with high pesticide uses. For each scenario, we held other predictors (nesting availability, abundance of *Protocalliphora* spp. within the nest, mean temperature, mean precipitation, and timing of clutch) at their population-level mean to isolate the direct effect of the agricultural landscape variables on body size. We generated 3,600 posterior draws from the fitted size model (A) using the *predicted_draws()* function to obtain population-level predictions, excluding random effects at the farm and nest-box levels. The resulting posterior predictive densities from both scenarios were visualized to compare puparial size density distribution.

## Results

### Variation in puparial size

Using a dataset of 1,513 measured *P. sialia* puparia collected between 2004 and 2019 (excluding 2008-2009) across a gradient of agricultural contexts, we observed notable variations in puparial area (mm^2^). The average puparial area was 26.85mm^2^ ± 3.47 SD, ranging from 10.74 mm^2^ to 37.80 mm^2^. These variations were analyzed using Model A (Table 1), which yielded a marginal *R^2^* of 17.2% and a conditional *R*^2^ of 37.6%, indicating that differences between years, farms, and nestboxes accounted for substantial variation.

Nestling availability (a proxy combining the number of living nestlings in a clutch and their duration availability for parasites) was positively correlated with puparial size, as clutches with few or short-lived nestlings consistently led to smaller puparia compared to larger or longer-lived clutches (Fig. 2a). Parasite density did not cause variation in puparial size, except in cases where there were more than 50 conspecific and/or heterospecific competitors inside the nest, although the trend seemed outlier-driven (Fig. 2b). Such cases (which occurred in only two nests in our sample), puparial size increased, whereas variance in individual size decreased, with almost no individuals being smaller than 22 mm^2^. The timing of *P. sialia* oviposition, estimated by the first hatching date of the Tree Swallow clutch (indicating when *Protocalliphora* larvae could begin feeding), showed a strong correlation with puparial size (Fig. 2c). Puparia in clutches initiated around day of year (DOY) 150 (end of May), averaged 27.5 mm^2^. In contrast, later clutches starting near DOY 180 (end of June) resulted in smaller puparia, averaging 25.2 mm^2^. Higher mean temperatures and precipitation during *Protocalliphora* spp. development were strongly and positively correlated with increased *P. sialia* puparial size (Fig. 2d,e). Predicted puparial area increased from 24.8 mm^2^ at 15°C to 28.2 mm^2^ at 22°C and from 25.2 mm^2^ to 27 mm^2^ at 1 to 7 mm of precipitation.

**Figure 2.**
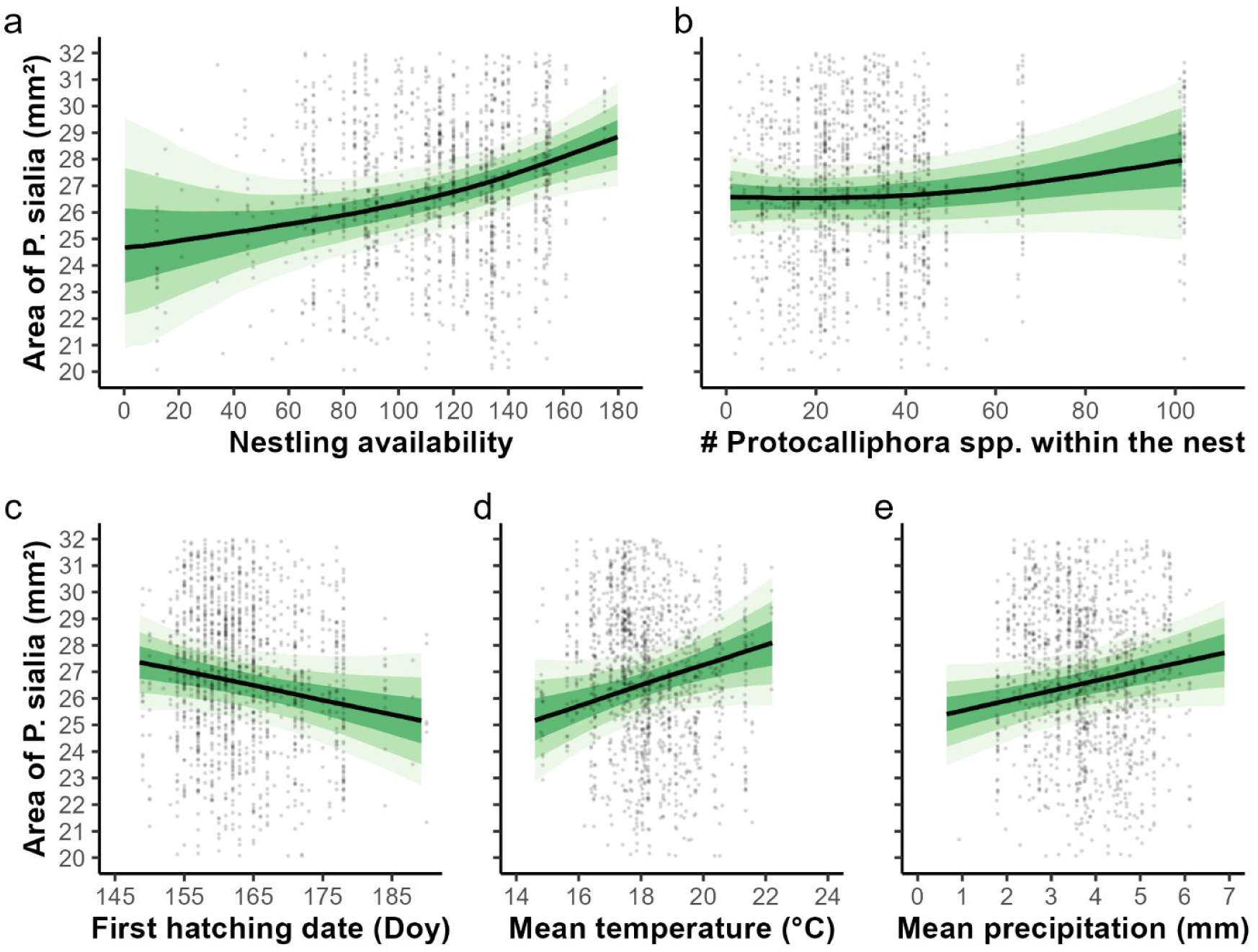
Variation in *P. sialia’s* puparia area (mm^2^) according to a) Nestling availability for their ectoparasites, b) Density of *Protocalliphora* spp. within the nest of measured *P. sialia*, c) First hatching date of the clutch from each nestbox, d) Mean temperature (°C) during *Protocalliphora* larvae development, e) precipitation (mm) during *Protocalliphora* larvae development. Grey dots indicate raw data. Credibility intervals (50%, 80% and 95%) are displayed in green.

Agricultural contexts surrounding the nests were characterized using a Robust Compositional Principal Components Analysis (rPCA; Fig. 1b). Component 1, which represents an agrointensive gradient, exhibited a non-linear effect on puparial size (Fig. 3a): mean area decreased from agroextensive (Scenario S1) to mid-intensity contexts, before increasing in highly intensive contexts (Scenario S2). These shifts were modest, ranging from 26.4 mm^2^ to 27.9 mm^2^. Conversely, puparial size variation (Fig. 3a) was lower in the agrointensive context (σ = 2.31) compared to the agroextensive contex (σ = 3.30), where individuals were, on average, smaller. Posterior predictive density distributions (comprising 3,600 random draws) confirmed that, all else being equal, sampling within an agrointensive landscape (S2) reveals an underrepresentation of small individuals and a narrower, right-shifted bell curve compared to agroextensive landscapes (S1; Fig. 3c). This suggests an underrepresentation of smaller phenotypes in agrointensive contexts.

**Figure 3.**
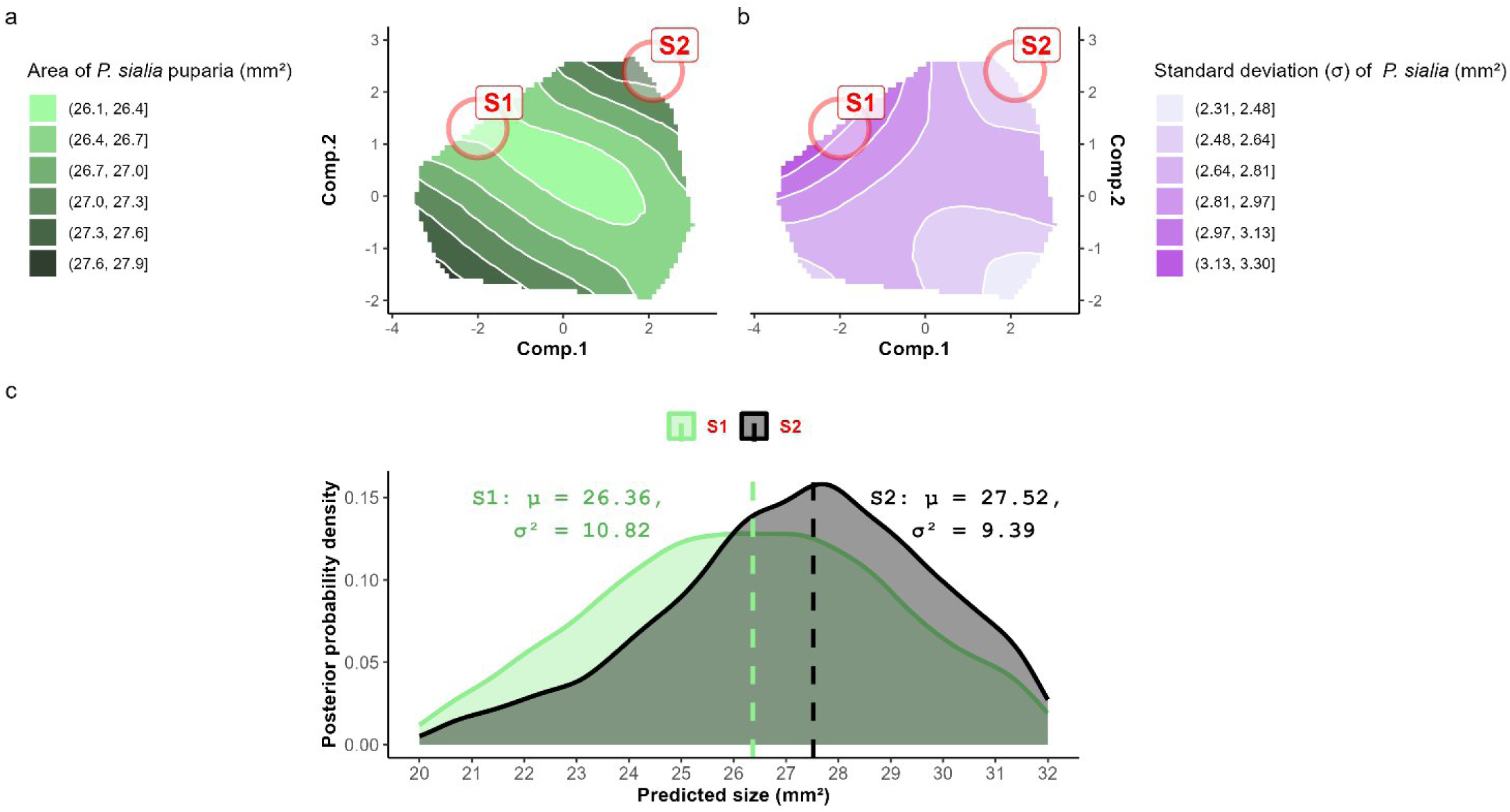
a) Heatmap displaying posterior predicted mean puparial size (mm^2^) of *P. sialia* across the agricultural gradient (Comp.1 and Comp.2 axes). Warmer colors indicate larger predicted body sizes. b) Heatmap of posterior predicted standard-deviation (σ) of *P. sialia* body size across the agricultural landscape. Warmer colors indicate higher heterogeneity in puparial size in different agricultural contexts. c) Posterior predictive density distributions of *P. sialia* puparial size under two contrasting agricultural scenarios: Scenario S1 (green) representing an agroextensive landscapes with high tree cover (Comp.1 = -2, Comp.2 = 1.3), and Scenario S2 (black) representing monoculture agrointensive contexts (Comp.1 = 2.3, Comp.2 = 2.4). Dashed vertical lines indicate posterior mean predictions for each scenario. Statistical parameters (μ and σ^2^) are displayed for each distribution.

### Influence of *P. sialia* morphological traits on infestation and emergence probabilities

Models B and C (Table 1) incorporate *P. sialia* puparial area (mm^2^) as an explanatory variable for both infestation and emergence probabilities using infestation and emergence data from 1,511 and 305 *P. sialia* puparia respectively, obtained marginal R^2^ values of 7.3% (conditional R^2^ = 17.6%) and 13.9% (conditional R^2^ = 29.5%). This highlights substantial variation attributed to temporal (year), spatial (farm), and nestbox-specific factors. Other diverse agroenvironmental predictors (*i.e. : agricultural landscape, weather, nestling host availability, Protocalliphora spp. density*) yielded results consistent with previous findings (see Coroller-Chouraki *et al*. 2026b) and can be find in Fig. S3.

The likelihood of *P. sialia* being infested by *Nasonia* wasps in our study system is high, with infestation rates exceeding 90% of *Protocalliphora* spp. individuals in some years (Coroller-Chouraki *et al*. 2026a). Puparial size had only a minor positive influence on this infestation probability. For instance, individuals with an area of 20.5 mm^2^ had an 81% probability of infestation compared to 88% for larger puparia measuring 32 mm^2^ (Fig. 4a). Similarly, emergence success (for puparia not infested by *Nasonia* spp.) was moderately influenced by size. Predicted emergence probability increased from 25% for smaller puparia (20.5 mm^2^) to 35% for larger ones (32 mm^2^), though these estimates were accompanied by wide credibility intervals, potentially caused by the smaller sampling size (Fig. 4b). The effects of other agroenvironmental parameters on emergence and infestation probability can be found in Fig. S3.

**Figure 4.**
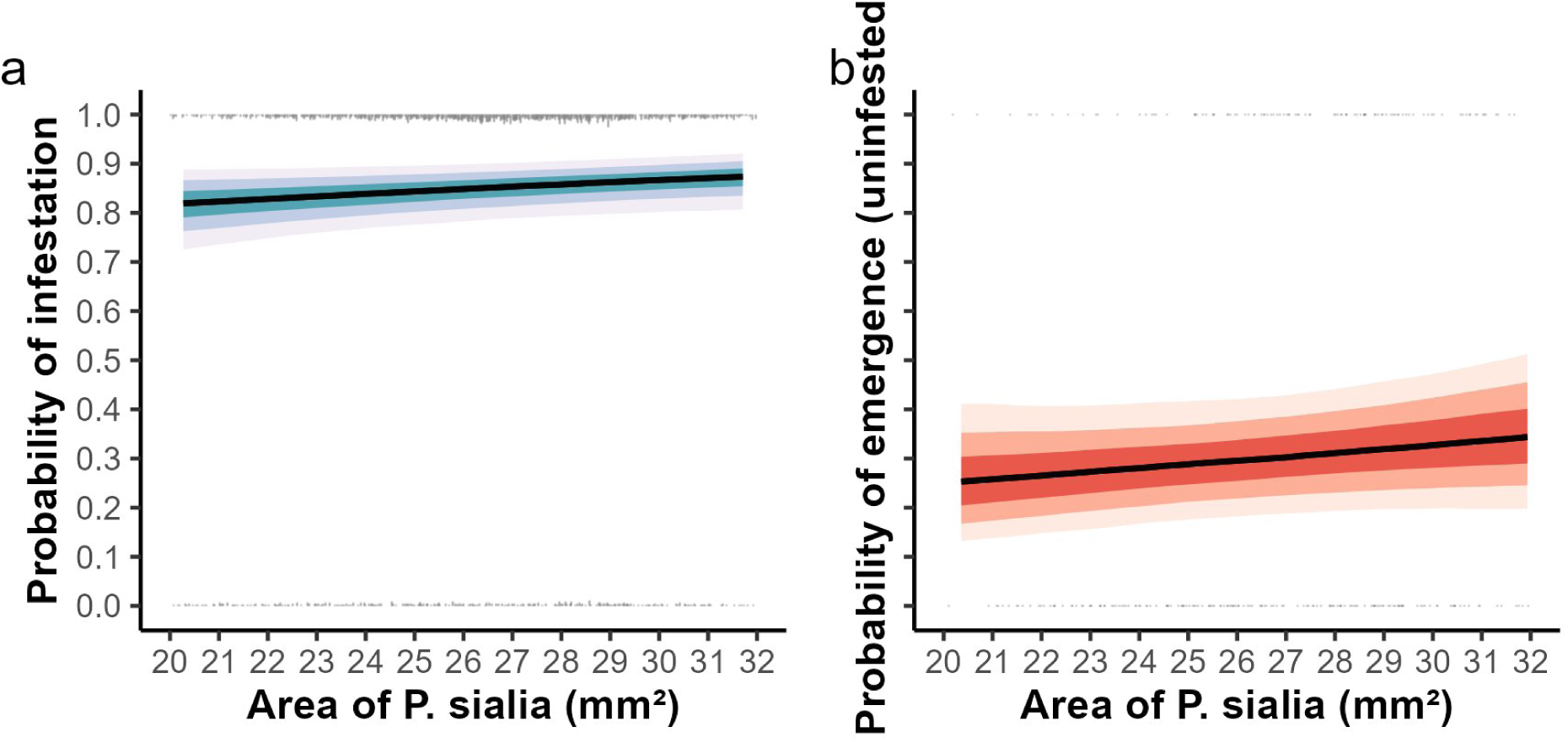
Influence of *P. sialia* size (area; mm^2^) on a) the probability of infestation by *Nasonia* parasitoid wasps (n = 1,511) and b) the probability of adult fly emergence for puparia not infested by *Nasonia* spp. (n = 305). Credibility intervals (50%, 80% and 95%) are displayed in green.

## Discussion

We found that *P. sialia* morphology is modulated by a combination of landscape composition, host resources, and ambient weather during the breeding phenology. However, this morphological plasticity has negligible consequences for both blowfly emergence and *Nasonia* parasitism. This lack of effect indicates that agricultural environments shape ectoparasite populations through direct extrinsic factors (e.g., thermal stress or landscape simplification). Crucially, these pressures are not mediated by phenotypic traits; a change in puparial size does not translate into altered life-history outcomes or shifted interactions with higher trophic levels

### Variation in *P. sialia* size

While density-dependent competition in host-parasite networks often shapes fitness through resource limitation for parasites and/or virulence for the host (Alizon and Lion 2011, Ben-Ami *et al*. 2008, Luong *et al*. 2011), our results suggest that both inter- and intra-specific competition were minimal in our system. Contrary to our expectation that high parasite density and low host availability would intensify scramble competition (Hughes *et al*. 2004), puparial size did not vary according to their interactions. Instead, it increased independently with host availability, suggesting that prolonged blood access enhances both growth and emergence success, consistent with known allometric relationships (Wainwright and Reilly 1994). Although food availability has been found to partially dictate adult size in Chown and Gaston (2010), Lambert (2007), and Nicol and Mackauer (2003), it appears that low host resources were not limiting at very high parasite densities in our study.

The slight (and possibly outlier-driven) increase in *P. sialia* puparial size at higher *Protocalliphora* spp. densities suggests several non-exclusive mechanisms. For example, a maternal effect, where higher-quality *Protocalliphora* females could produce more and fitter offspring. Alternatively, high *Protocalliphora* densities could improve group feeding efficiency (Peters and Barbosa 1977). Finally, the selective mortality of smaller individuals before pupation and sampling could artificially inflate the observed mean size.

Seasonality and phenology also emerged as important drivers of size as *P. sialia* puparia developing on later Tree Swallow clutches reached smaller puparial sizes than those from earlier clutches. This suggests a *P. sialia* egg quality decrease over time (Cağlar *et al*. 2014, Walters and Hassall 2006) or a preferential allocation of high-quality eggs to earlier clutches, in which nestling host availability is higher (Coroller-Chouraki *et al*. 2026b). Additionally, late-season nestlings are often in poorer condition (Harriman *et al*. 2017), likely providing lower-quality blood meals that constrain larval development and highly limit emergence success (Coroller-Chouraki *et al*. 2026c). Finally, the positive correlation between *P. sialia* puparial size with both precipitation and temperature mirrors our previous observations where *Protocalliphora* spp. were more abundant in nests exposed to similar weather conditions (Coroller-Chouraki *et al*. 2026b).

*Protocalliphora sialia* puparia were slightly larger in two contrasting landscapes: agrointensive areas (monocultures with high pesticide use) and agroextensive pastures. Conversely, body size decreased in the most *Protocalliphora*-abundant habitats, typically agroextensive areas with high tree cover (Coroller-Chouraki *et al*. 2026b). Rather than suggesting that intensive farming enhances individual growth, these patterns likely reflect survival and sampling biases. Beyond reduced population densities (Coroller-Chouraki *et al*. 2026b), agricultural intensification appears to selectively truncate *Protocalliphora* size distributions. Our finding that *P. sialia* distributions are more left-skewed in intensive landscapes (combined with the inherent scarcity of small-sized outliers, Coroller-Chouraki *et al*. 2026a) suggests that in harsh agrointensive settings, stressors such as pesticides and landscape homogeneity likely exert selective survival pressure. This biological filtering could remove small phenotypes from the sampled population, resulting in a truncated size distribution with a higher mean and a lower variance. Conversely, favorable agroextensive landscapes would allow a broader range of phenotypes, including smaller individuals, to survive until pupation.

Impact of size on *P. sialia* emergence and host-selection by *Nasonia* spp.

Our models reveal that *P. sialia* puparial size is highly plastic, driven by host availability, weather, and agricultural context. The cumulative effect of these factors is substantial; while host availability alone induced a 16% change in mean area (25 to 29 mm^2^), the full range of observed sizes spanned 20 to 32 mm^2^. Despite this variability, puparial size explained less than 10% of the variation in emergence success or *Nasonia* spp. parasitism. Several factors likely account for this weak relationship.

First, *Nasonia* infestation rates in our system were consistently high, averaging 68.8% and peaking at 90.5% (Coroller-Chouraki *et al*. 2026a) and far exceeding the 16.7% rate previously reported in North America (Bennett and Whitworth 1991). These patterns are likely to decrease the statistical power to detect size-dependent selection, as non-infested puparia were relatively rare. Furthermore, the high infestation rate suggests a generalist host-use strategy by *Nasonia* in this system, where oviposition may be driven more by host density and nest microclimate than by individual puparial size (Coroller-Chouraki *et al*. 2026b). While finer-scale decisions like egg allocation or sex ratio adjustment may still be influenced by host quality (Godfray 2010; Shuker et al. 2005), our *in-vivo* study suggests that host size might not be a gate keeper for selection in our system.

Second, while *in vitro* experiments show that *Nasonia* spp. adjust clutch size and sex ratio based on host quality, species, and prior parasitism (Koppik et al. 2014, 2019; Rivers & Denlinger 1995; Shuker et al. 2005), our *in vivo* study only assessed the binary outcome of host acceptance or rejection. To our knowledge, experimental evidence for *Nasonia* rejection behavior based solely on intraspecific puparial size variation is lacking, as rejection typically occurs between different host species (Rivers & Denlinger 1995).

Third, the survival and sampling biases inherent to our study system may have dampened observable fitness outcomes. If the smallest, most vulnerable larvae failed to reach pupation, our analysis was restricted to a truncated population of higher-quality survivors. This biological filtering combined with the limited sample of measured and uninfested *P. sialia* (n = 305), likely obscured the relationship between size and emergence success, an effect observed in other Diptera (Levot et al. 1979, Packer and Corbet 1989).

Previous results showed that *Protocalliphora* spp. emergence in unparasitized individuals was lower in forest-rich agroextensive landscapes where *Protocalliphora* were most abundant, compared to both agrointensive and agroextensive landscapes with less tree cover, where they were less numerous (Coroller-Chouraki *et al*. 2026c). This spatial variation in emergence tracks specific morphological shifts; habitats with lower parasite numbers are characterized by significantly larger individuals and reduced size variance, while high-density agroextensive environments preserve a broader, unfiltered phenotypic diversity. Together, these findings support the hypothesis that benign, forested environments allow a greater spectrum of phenotypes (including smaller, less competitive individuals) to pupate, reducing mean size. Conversely, harsher landscapes filter out weaker individuals prior to pupation, artificially inflating average size metrics. Our findings highlight the importance of accounting for missing fractions when interpreting trait variation such as puparial size. In agrointensive landscapes, increased mortality of smaller or lower-quality larvae prior to sampling likely biased our data toward larger individuals, masking potential bigger positive effects of size on emergence success or negative effects on host selection by *Nasonia* parasitoids. Had we been able to track larval survival throughout development, we might have observed stronger size-dependent effects. Such survival biases can obscure key life-history patterns, yet recognizing them is crucial for understanding population dynamics and anthropogenic impacts on trophic networks involving hosts, parasites and parasitoids.

## APPENDIX 1

**Figure S1.**
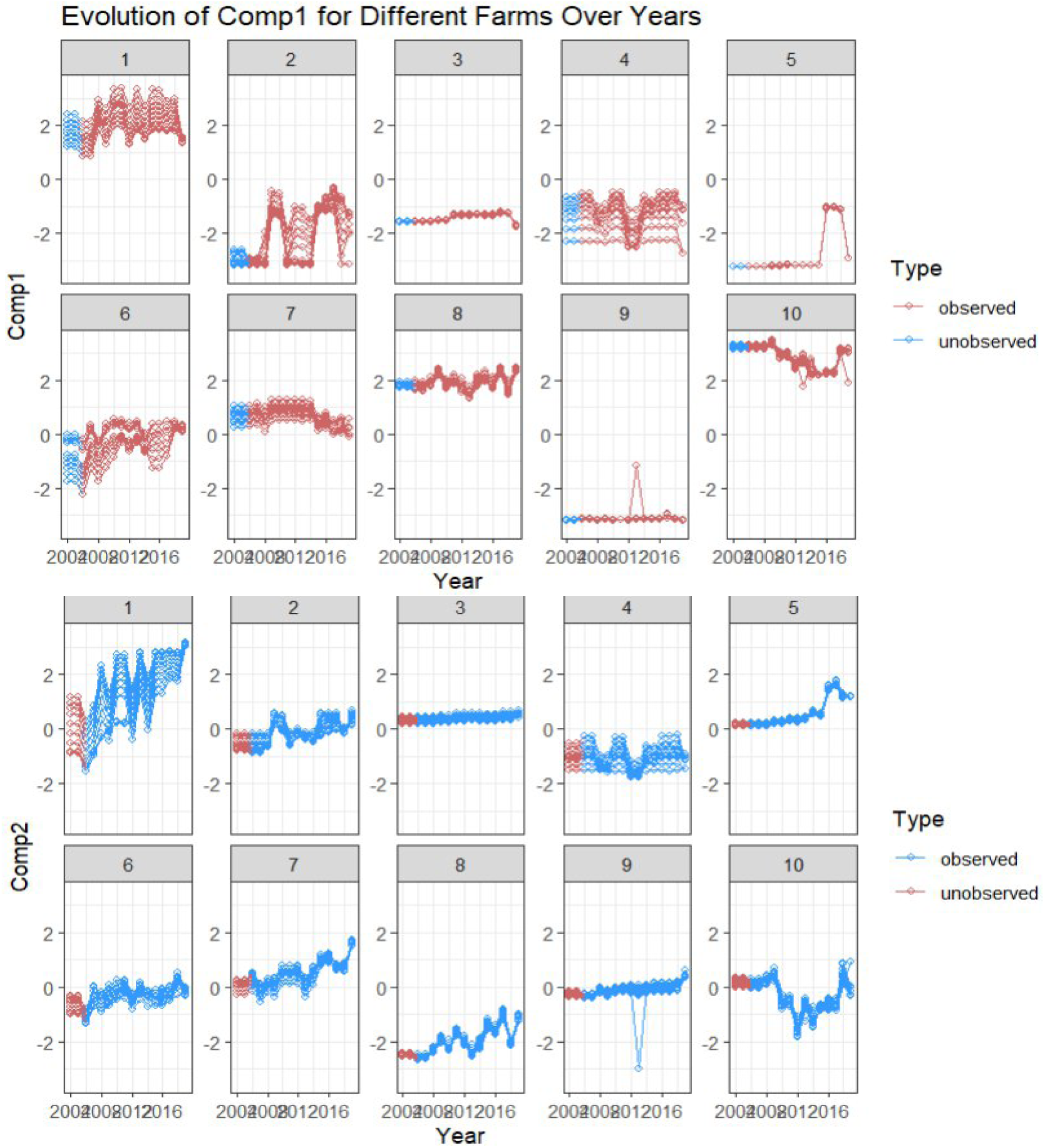
Interannual stability of landscape composition around Tree Swallow nest boxes between 2004 and 2019. Extrapolated Comp.1 or Comp.2 for 2004 and 2005 missing values are displayed in contrasting color (respectively blue vs red, and red vs blue).

### Directed Acyclic Graph (DAG) linking *Protocalliphora sialia’s* size to their emergence success

**Figure S.2.**
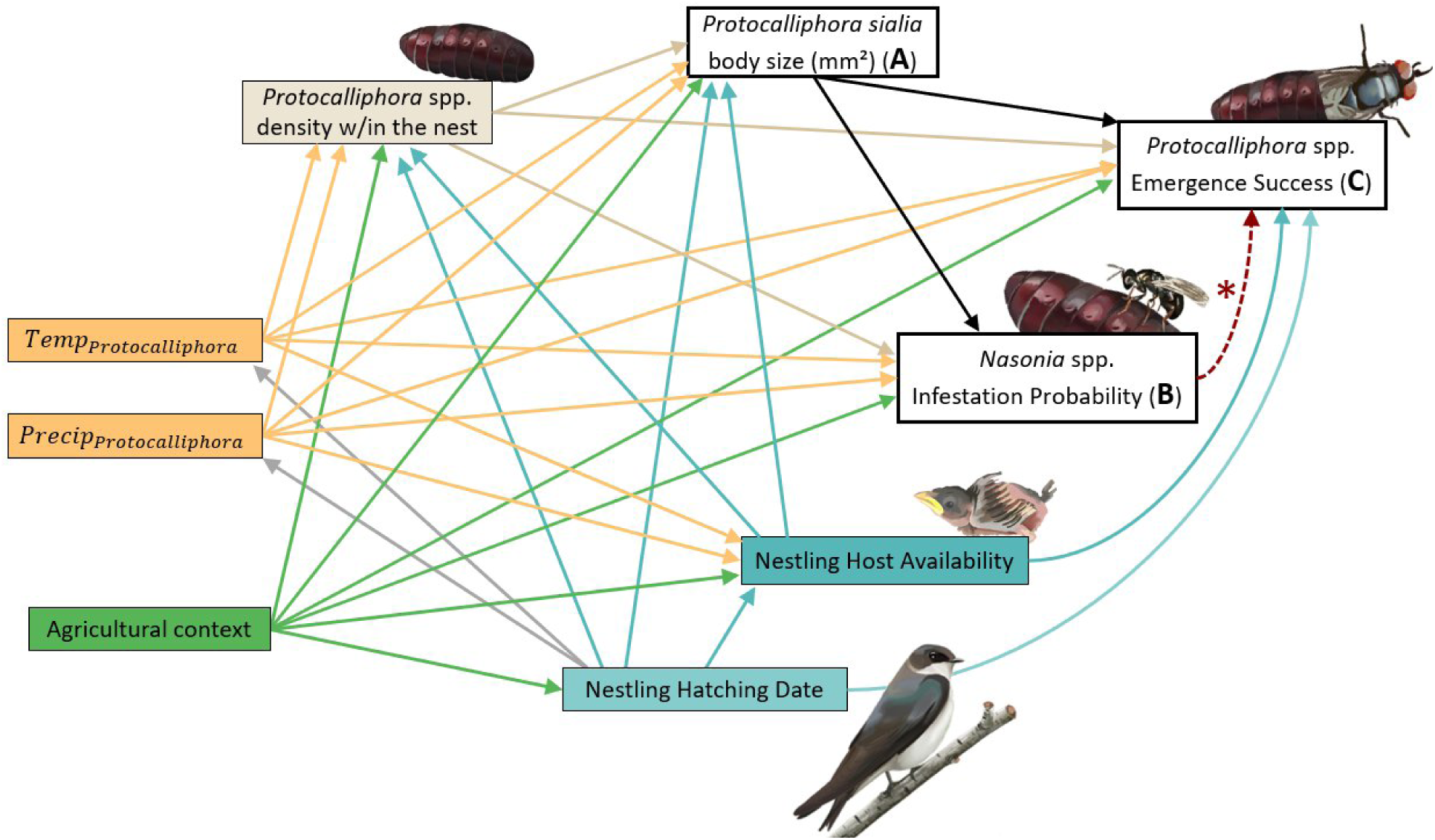
Directed Acyclic Graph of putative causal relationships among studied variables. Direct effects of variables on (A) *Protocalliphora sialia* body size (mm^2^), (B) the probability of *Nasonia* infestation in *P. sialia* puparia, and (C) the emergence probability in *Protocalliphora* puparia (see models in Table 1). Weather conditions during the larval/pupation phases corresponded to hourly averages over the respective period during which swallow nestlings and blowfly larvae and pupae developed in each swallow brood (i.e., up to fledging or death and to estimated emergence, respectively). Agricultural context surrounding each swallow brood was defined based on the scores of the first two axes of a robust, compositional PCA fitted on relative land cover values measured annually within 500 m of each nest box of the study system (i.e., Comp.1 and Comp.2 in Fig. 6B). * The red dotted arrow indicates a deterministic biological constraint: infestation by *Nasonia* spp. systematically leads to *Protocalliphora* mortality, resulting in zero emergence success (perfect separation). To identify the effects of body size on *Nasonia* infestation probability, all *P. sialia* puparia were included in a first model (A; Table 1). A second model was then performed strictly on uninfested *P. sialia* to assess the impact of body size and environmental drivers on emergence success (B; Table 1).

### Modelization of the relationship between Hatching Date (phenology) and mean temperature during nestling development

**Script S1**. Model output and prediction corresponding to the relation between Hatching Date (phenology) and mean temperature during nestling development

`summary(Modele_Protocalliphora_sialia_parasitoidism)`

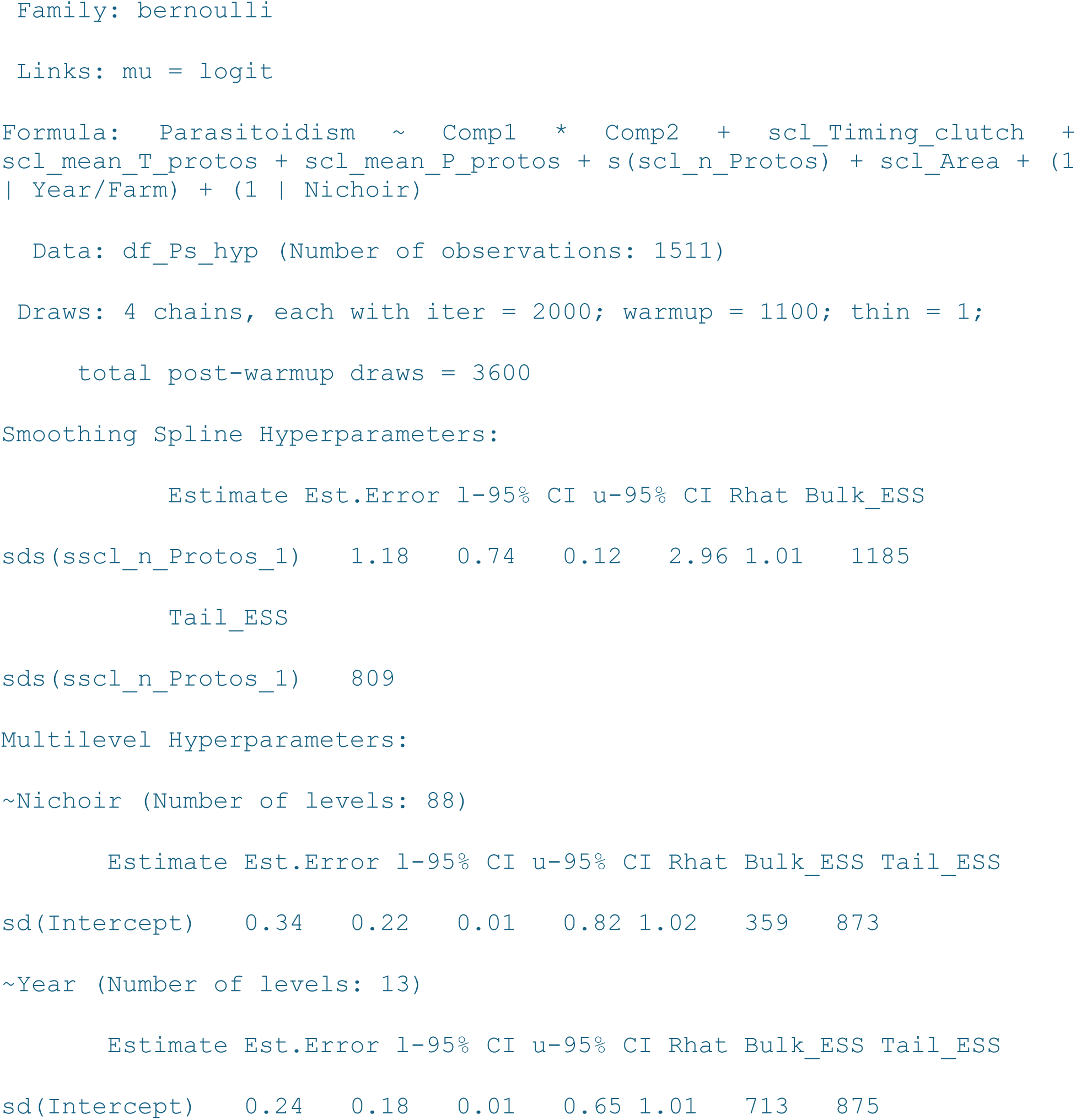
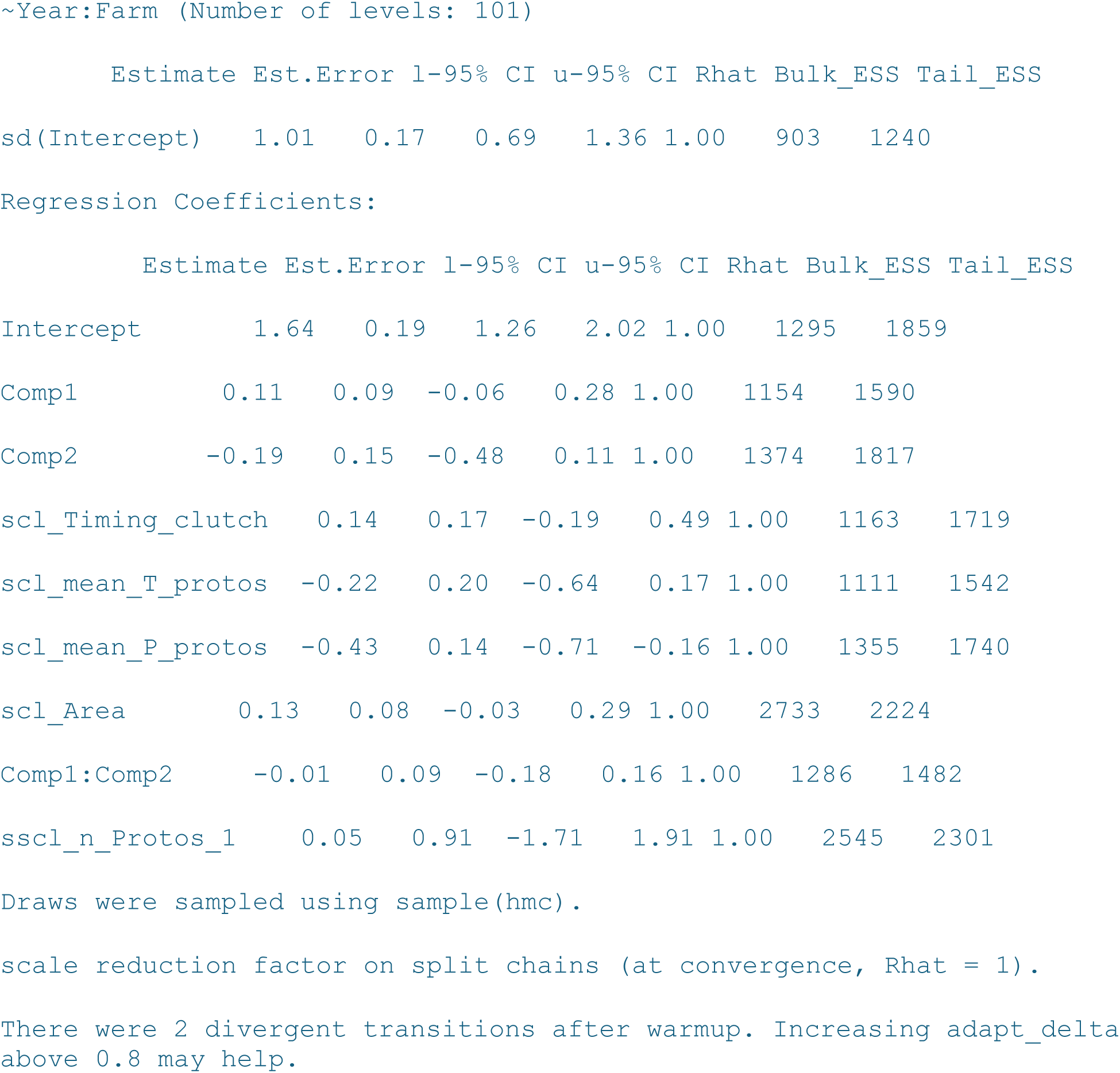

`Summary(Modele_Protocalliphora_sialia_emergence)`

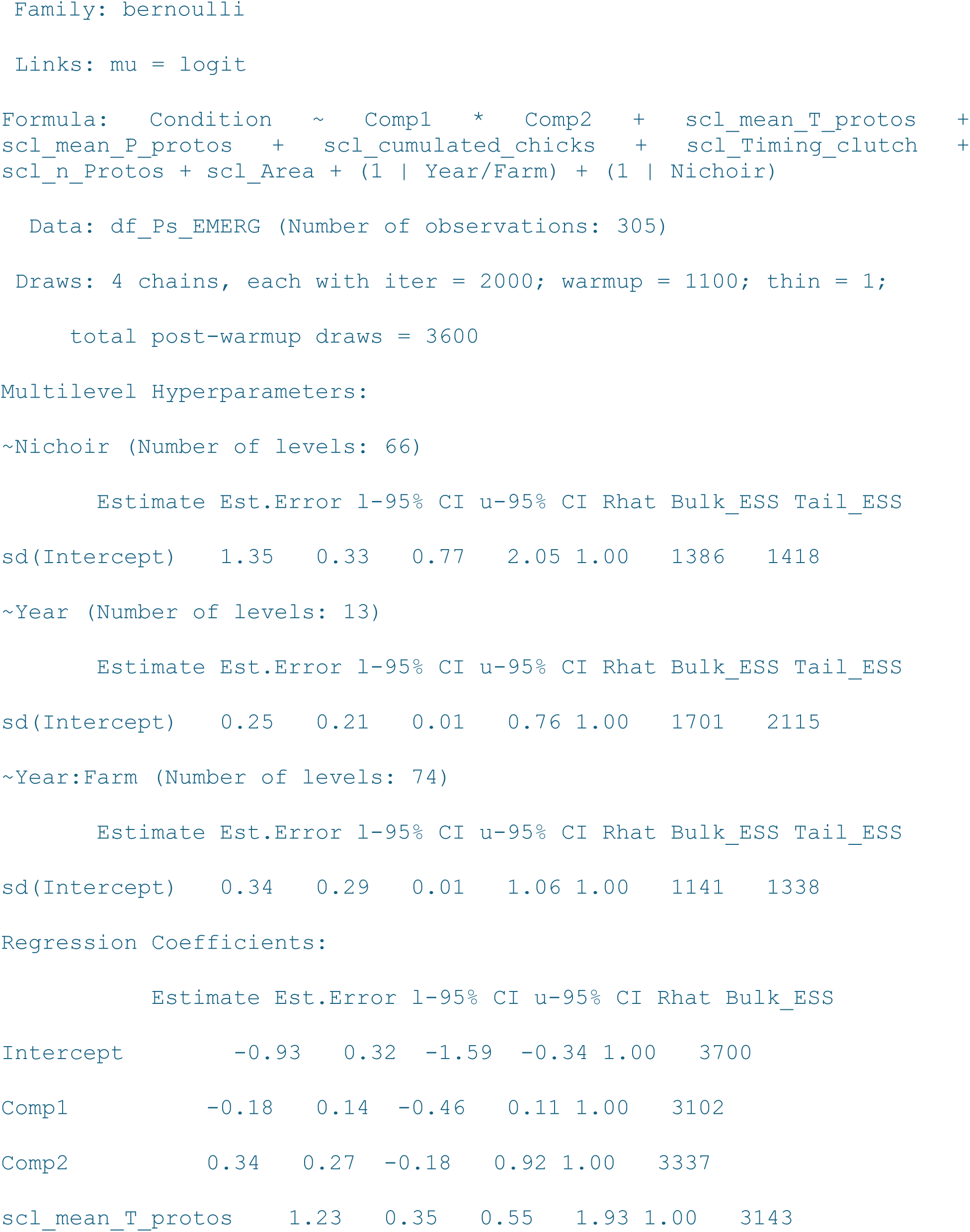
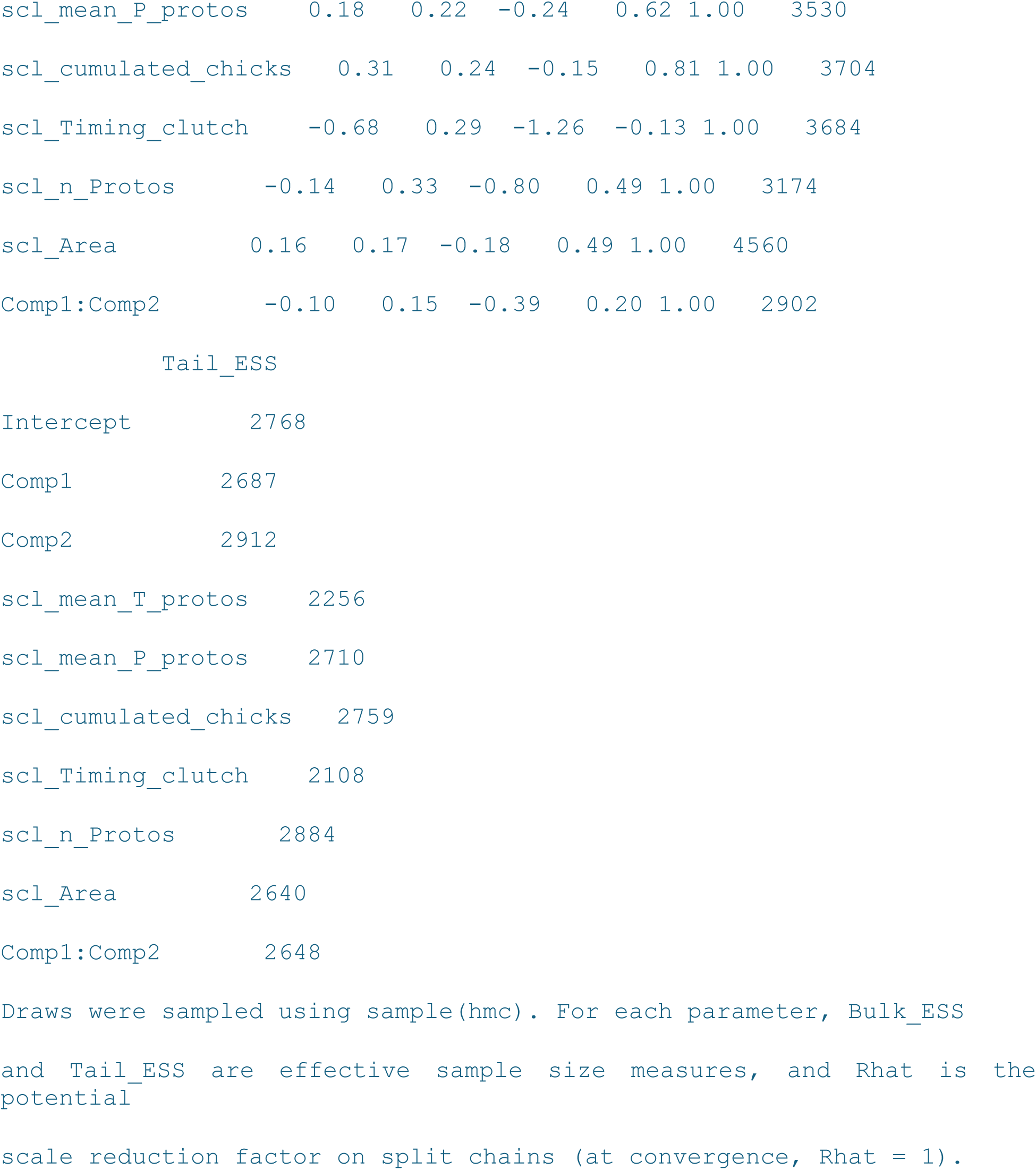

**Figure S3.**
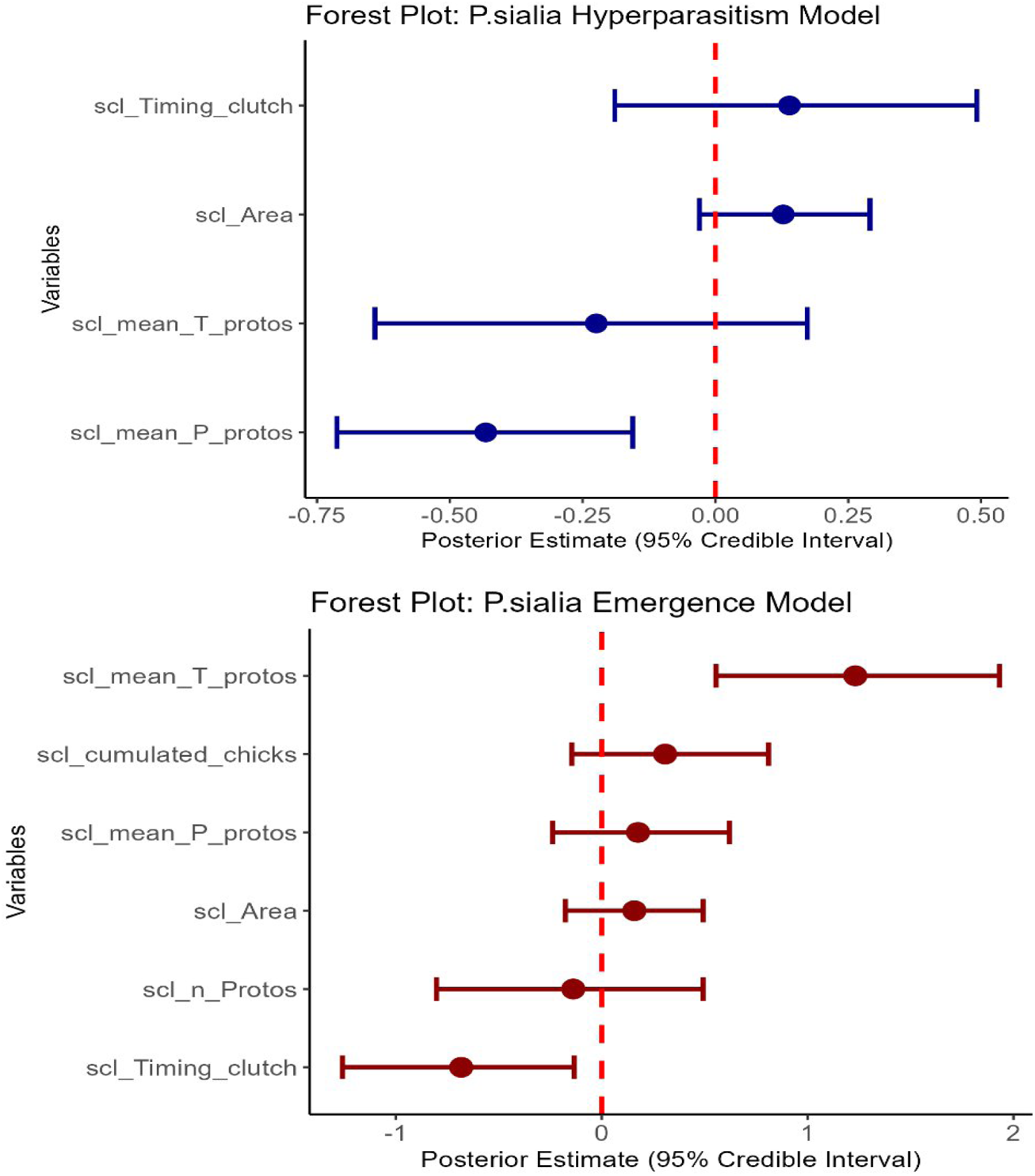
Posterior estimates (median ± 95% credibility interval) of main effects of several factors on *Protocalliphora sialia* parasitoidism risk or emergence probability. The vertical dashed red line at zero represents no effect. Effects are in log-odds; positive values increase parasitoidism or emergence probability. Splines and interactions are excluded from forest plot as a single coefficient doesn’t capture the full relationship of a spline (Wood, 2017*) and as the marginal effect of the constituent variables of an interaction cannot be examined in isolation (Brambor *et al*. 2006**). * Wood, S. N. (2017). *Generalized additive models: an introduction with R*. chapman and hall/CRC. ** Brambor, T., Clark, W. R., & Golder, M. (2006). Understanding interaction models: Improving empirical analyses. *Political analysis*, *14*(1), 63-8.

## Notes

### Competing Interest Statement

The authors have declared no competing interest.

